# The impact of *Bacillus coagulans* X3 on available nitrogen content, bacterial community composition, and nitrogen functional gene levels when composting cattle manure

**DOI:** 10.1101/2024.02.23.581714

**Authors:** Biao Liu, Wei Chen, Zhen Wang, Zhaohui Guo, Yongmei Li, Lijuan Xu, Minxi Wu, Hongmei Yin

## Abstract

This study was designed to probe available nitrogen levels, bacterial community composition, and the levels of nitrogen functional genes present when composting cattle manure with or without the addition of *Bacillus coagulans* X3. Bacterial supplementation was associated with the prolongation of the thermophilic stage and improved maturity of the resultant compost. At the maturity stage, samples to which *B. coagulans* X3 had been added exhibited significant increases in ammonium nitrogen, nitrate nitrogen, and total nitrogen levels. The dominant bacterial phyla observed in these composting samples were Firmicutes, Proteobacteria, Bacteroidetes, Actinobacteriota, and Chloroflexi*.B. coagulans* X3 addition resulted in significant increases in relative Firmicutes abundance during the thermophilic and cooling stages, while also increasing *amoA* and *nosZ* gene abundance and reducing *nirS* gene levels over the course of composting. Together, these data suggest that *B*. *coagulans* X3 supplementation provides an effective means of enhancing nitrogen content in the context of cattle manure composting through the regulation of nitrification and denitrification activity.

## 1. Introduction

Rapid advances in the development of large-scale poultry and livestock industries in China have given rise to high levels of animal manure output [1]. Feasible approaches to treating and utilizing this manure are essential in order to minimize the risk of environmental pollution. Aerobic composting has been established as a safe and effective approach to manure treatment, with the end products of such composting offering further value as nutrient-rich materials that can be safely employed as organic agricultural amendments [2-4]. The biochemical process of composting entails the microbe-mediated degradation of organic matter to produce humus-like byproducts[5,6]. Many challenges can arise when biodegradation activity is poor or there are insufficient microbes present in materials being composted, contributing to low organic matter degradation efficiency, prolonged fermentation cycles, and the loss of significant nitrogen [7,8]. A range of methods have been employed to date in an effort to improve manure composting quality and efficiency, such as the adjustment of moisture levels, the modulation of the C/N ratio, the optimization of ventilation, or the use of additives [9,10].

Exogenous microbe supplementation has emerged as a promising approach to reducing composting cycle length, minimizing nitrogen loss, and improving the overall quality of compost[11]. When seeking to improve nitrogen content and maturity in their study of pig manure composting, Jiang et al. [12] found that there were benefits to adding a 1% nitrogen turnover bacterial agent consisting of a range of ammonifiers, nitrobacteria, and *Azotobacter* species at the start of the compositing process. Mao et al. [13] further determined that total nitrogen (TN) and dissolved organic carbon levels in composted pig manure could be enhanced by bacterial addition. Li et al. [14] determined that microbial inoculation (*Acinetobacter pittii*, *Bacillus subtilis* subsp. *Stercoris*, and *B. altitudinis*) was sufficient to prolong the thermophilic composting stage while increasing overall high-temperature-resistant bacterial abundance and ammonium nitrogen(NH_4_^+^-N) and nitrate nitrogen(NO_3_^—^N) concentrations in composted materials. Analyses of metabolic function have provided further evidence for the ability of microbial inoculation to reduce the levels of human disease-related functional genes while increasing carbohydrate metabolism levels.

Exogenous microbial inoculation has also been shown to have a significant impact on the structure of microbial communities within compost samples. For example, Li et al. [14] observed that such inoculation led to enhanced Moraxellaceae proliferation on day 4 of composting, together with an increase in Planococcaceae abundance on days 12-24 of composting. Guo et al.[15] explored the impact of adding *Bacillus megaterium* on the dynamics of microbial communities in pig manure undergoing aerobic composting, revealing that these additives resulted in a 48.05% increase in Firmicutes abundance on day 17, while there was a 30.89% reduction in Proteobacteria abundance in the final compost products as compared to control conditions.

A series of nitrogen functional genes are responsible for the transformation of nitrogen during composting, including the processes of nitrogen mineralization, ammonification, nitrification, denitrification, and nitrogen fixation[16]. Yang et al. [17] found that microbial activity and nitrogen functional gene expression levels were closely associated with changing nitrogen content levels. Ammonia nitrogen oxidation to nitrate nitrogen is primarily regulated by ammonia monooxygenase (*amo*A), while NO_2_^-^ conversion to NO is under the control of denitrifying microbes that express nitrite reductase (*nir*K and *nir*S) genes. The enzyme encoded by the nitrous oxide reductase (*nosZ*) gene is responsible for converting N_2_O to N_2_ [18]. In several reports, exogenous microbe inoculation was found to alter nitrogen functional gene abundance when composting solid waste, thereby influencing nitrogen transformation [15,19].

In a previous study, our group isolated the thermotolerant keratin-degrading *Bacillus coagulans* X3 strain from soil samples [20]. This strain is capable of producing cellulases and proteases such that it is a promising tool for use in the context of organic waste degradation. The specific effects of *B. coagulans* X3 inoculation on nitrogen transformation in the context of cattle manure composting, however, remain to be assessed. Accordingly, this study was designed to examine the impact of such microbial inoculation on available nitrogen content, bacterial community characteristics,and nitrogen functional genes when composting cattle manure.

## 2. Materials and Methods

### 2.1 Composting materials, study design, and sample collection

Fresh cattle manure and rice straw were obtained from a cattle farm and local residents in Yiyang, Hunan Province, China. Rice straw was air-dried, after which it was cut into ∼3-5 cm lengths. The physicochemical properties of the raw materials used for this study are presented in Table 1. Prior to composting, *B. coagulans* X3 was activated and expanded in culture, with cultures then being centrifuged, washed, and resuspended usingsterile distilled water. Resuspended bacteria were adjusted to a concentration of ∼10^9^ CFU/mL for subsequent utilization.

**Table 1.** Raw material physicochemical properties.

| Parameters | Cattle manure | Rice straw |
| --- | --- | --- |
| Moisture(%) | 61.22±0.75 | 9.78±0.38 |
| pH | 8.23±0.03 | 7.00±0.02 |
| Total carbon(%) | 33.55±0.34 | 42.16±0.48 |
| Total nitrogen(%) | 1.99±0.03 | 0.71±0.02 |
| C/N ratio | 16.89±0.10 | 59.67±0.65 |

All composting experiments were performed at Yiyang Yuanfeng Biotechnology Co., Ltd, China over a 35-day period. Briefly, cattle manure and rice straw were adjusted at a 5:1 (w/w) ratio to adjust the C:N ratio of the resultant compost to 25:1. Moisture content in the compost was then adjusted to ∼60%. Two composting treatment conditions were established, each using a composting pile that was 3.0 m long × 2.0 m wide × 1.5 m tall. For the test group (designated T),1% *B. coagulans* (inoculant volume/wet composting sample weight) was added, while no inoculation was performed for the control group (designated CK). Three replicates were established for each treatment. To ensure the greatest efficiency of the composting fermentation reaction and to ensure that sufficient oxygen was available, The compost pile was turned every 3 days during the first 15 days and every 5 days thereafter.

On days 0, 3, 6, 9, 15, 25, and 35, compost samples were collected. Prior to the collection of these samples, all compost piles were turned to ensure uniformity. Equal sample amounts were collected at random from all compost piles using a five-point sampling approach. After samples had been mixed to ensure that sampling was performed in a representative manner, ∼500 g of each sample (fresh weight) was gathered. These samples were then separated into three parts, with one being freeze-dried and stored at -80°C for subsequent 16S rDNA sequencing and analyses of nitrogen functional genes, one being stored at 4°Cto analyze the germination index(GI) analysis, and one being air-dried to analyze pH, TN, NH_4_^+^-N, and NO_3_^-^-N levels.

### 2.2 Physicochemical analyses

Adigital thermometer (Yidu, LCD-105,China) was used to assess compost temperatures at 9:00 and 15:00 each day, with average temperatures being recorded. Average ambient air temperatures on a given were calculated based on the environmental temperature. To measure pH values, 5 g of air-dried samples were added to 50 mL of deionized water for 30 min and shaken, followed by analysis with a pH electrode (INESA, PHS-3E, China). To extract NH_4_^+^-N and NO_3_^-^-N, 2 M KCl (1:20) was used, and analyses were conducted using a colorimetric approach [21].TN levels and GI values were assessed as per the Chinese standard for organic fertilizer(NY525-2021).

### 2.3 High-throughput 16S rDNA sequencing

Based on measures of composting temperatures, freeze-dried samples collected on days 3 (mesophilic stage), 6 (thermophilic stage), 15 (cooling stage), and 35(maturation stage) were selected for high-throughput 16S rDNA sequencing performed by Shanghai Majorbio Biotechnology Co.Ltd(Shanghai,China). Briefly, an E.Z.N.A soil DNA kit (Omega Biotek,GA, USA) was used to extract total genomic DNA, after which the purity and concentration of these extracted DNA samples were measured via1% agarose gel electrophoresis and through the use of a Nanodrop 2000 instrument. The 16S rDNA V3-V4 hypervariable region was amplified with the 338F (5′-ACTCCTACGGGAGGCAGCAG-3′) and 806R (5′-GGACTACHVGGGTWTCTAAT-3′) PCR primers using the following thermocycler settings: 95°C for3 min; 30 cycles of 95°C for 30 s, 55°C for 30 s, 72°C for45 s; 72°C for 5 min. PCR products were analyzed and purified via 2% agarose gel electrophoresis using an AxyPrep DNA Gel Extraction Kit (Axygen Biosciences,USA), recovering target bands [22]. The Majorbio Cloud Platform was used to analyze data generated with the Illumina MiSeq PE300 platform.

### 2.4 qPCR

Nitrogen functional gene(*amoA*,*nirK*,*nirS,*and *nosZ*) abundance in different samples was quantified via qPCR using the primers presented in Table 2, which have also been reported previously[17]. All qPCR analyses were performed with an ABI 7300 real-time PCR instrument using 20μL reactions containing 10 μL of 2× ChamQ SYBR Color qPCR Master Mix, 0.8 μL of each primer, 6 μL of ddH_2_O and 2 μL of templateDNA. Thermocycler settings were: 95°C for 3 min; 40 cycles of 95°C for 5 s, 58°C for 30 s, and 72°C for 60 s, with a final melt curve analysis. Quantification of all genes was performed in triplicate using a standard curve prepared from serial 10-fold dilutions of plasmids containing the cloned *amo*A, *nir*S, *nir*K, and *nos*Z genes. The respective amplification efficiencies for these genes were 100.65%, 93.09%, 101.30%,and 103.72%.

**Table 2.**
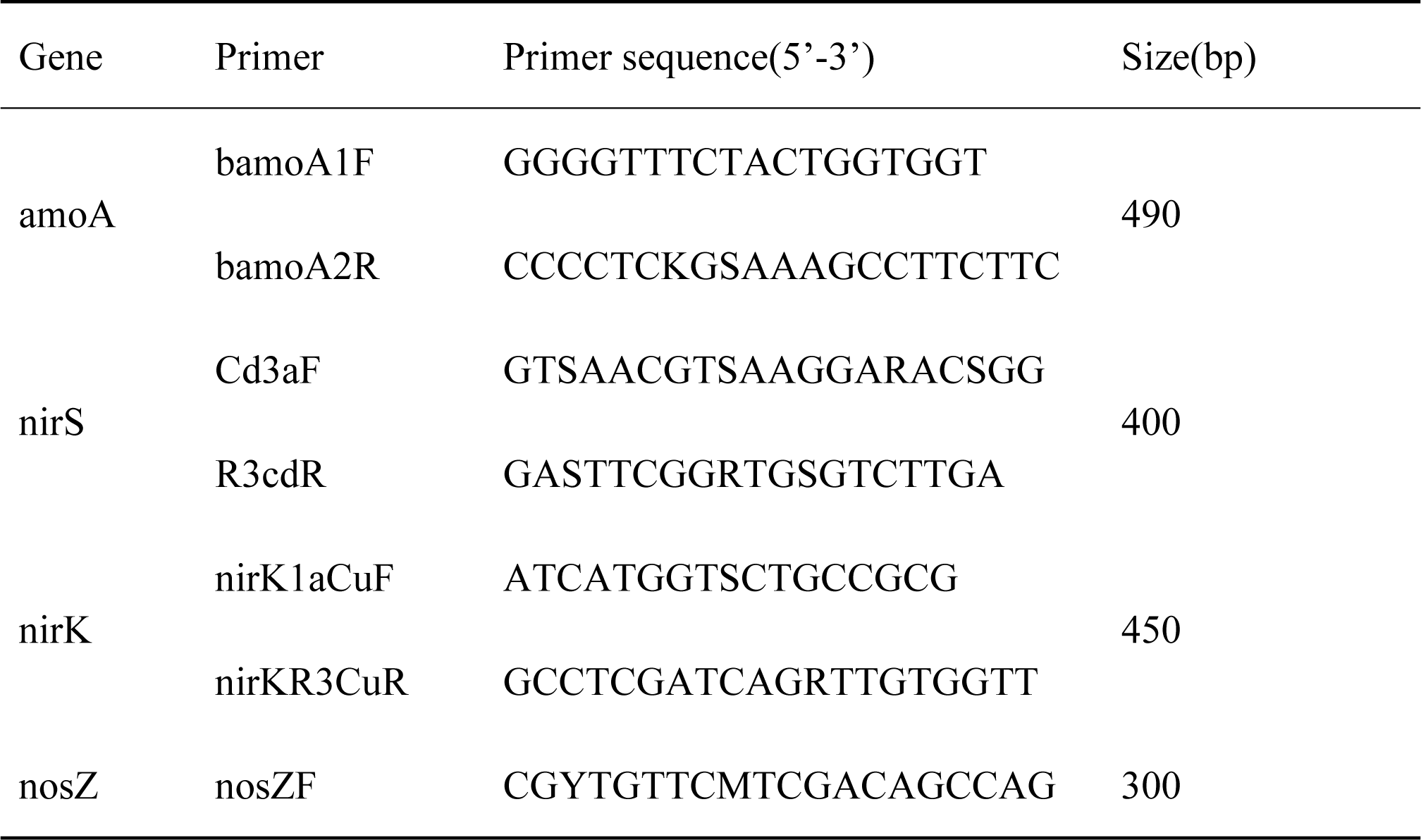
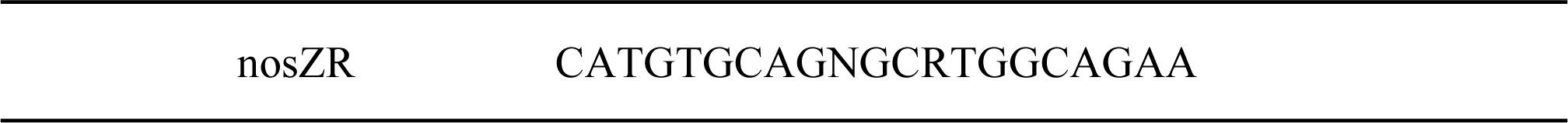
Primer sequences used for this study.

### 2.5 Data analyses

Data are given as means± standard deviation from experiments performed in triplicate. Microsoft Excel and SPSS 25.0 were used to analyze all data related to physicochemical properties and nitrogen functional gene abundance. Bacterial community analyses were conducted with the online Majorbio Cloud Platform. Raw sequences were submittedto the NCBI Sequence Read Archive database (PRJNA916590).

## 3. Results and Discussion

### 3.1 Changes in temperature, pH, and GI during composting

Compost temperatures offer valuable insight into the intensity of microbial activity and the harmlessness of the composted products[23].Both the CK and T treatment groups exhibited a typical trend in temperatures over the course of composting consisting of mesophilic, thermophilic, and cooling-maturation stages. The temperature in the CK group of 50°C was reached on day 3 and remained above this level for 9 days, reaching a maximum of 58.5°C. In group T, a temperature of 50°C was recorded on day 2, and it remained above this level for 12 days, peaking at 61.9°C (Fig. 1a). Thermophilic stages of these durations were sufficient to kill weed seeds and potential pathogenic microbes, thereby allowing the resultant compost samples to meet the Chinese standard for Technical specification for sanitation treatment of livestock and poultry manure (GB/T 36195-2018). These results support the ability of *B. coagulans* X3 inoculation to prolong the thermophilic stage during composting and to increase the temperature of the composting reaction. This is consistent with similar results reported previously following ABB consortium (*Acinetobacter pittii*, *Bacillus subtilis* subsp. *Stercoris*, and *Bacillus altitudinis*) [14] or *Aneurinibacillus* sp. LD3[24] inoculation during composting.

**Fig. 1.**
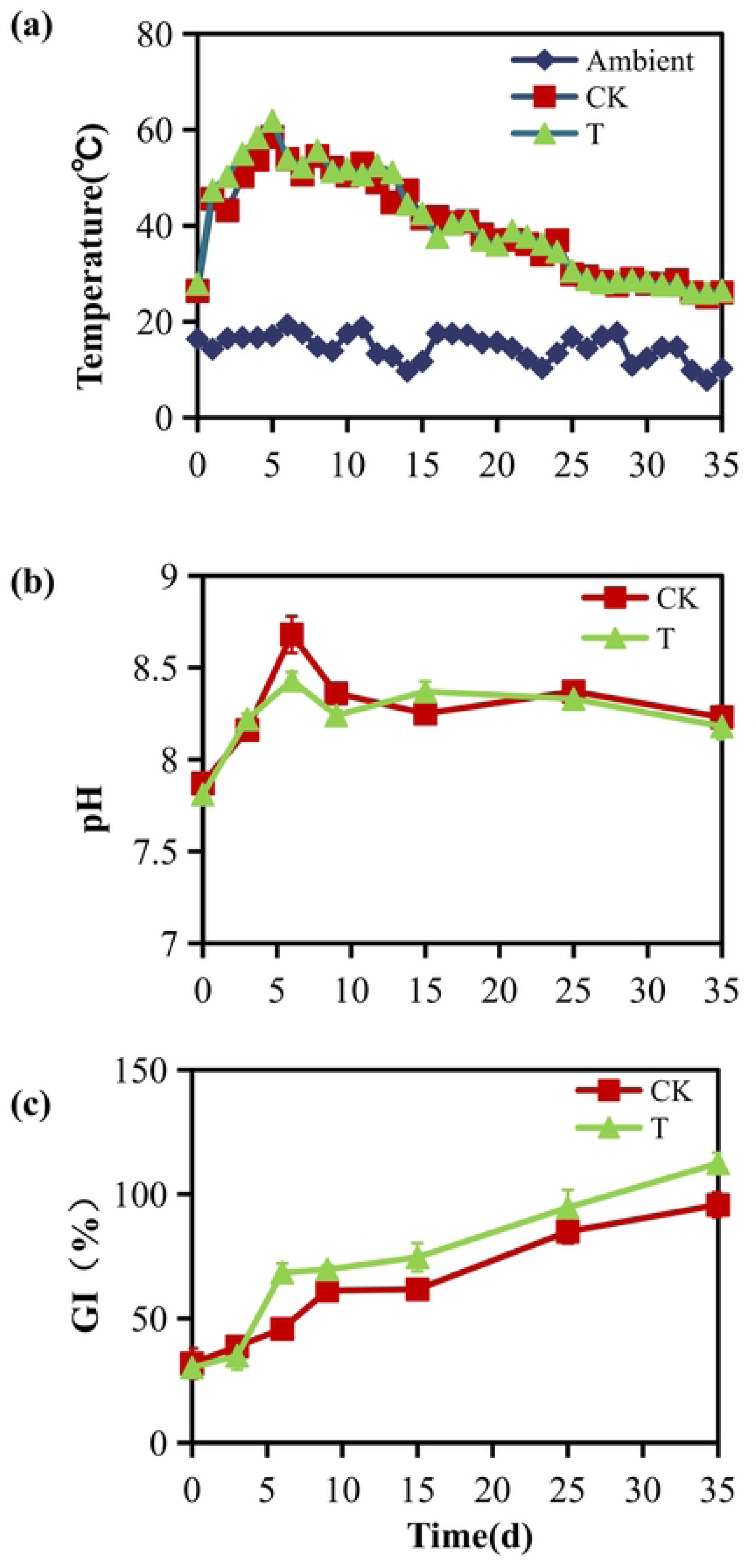
Temperature(a), pH(b), and GI(c)changes in the indicated groups over the course of composting(CK, no additive; T, 1% *Bacillus coagulans* X3).

As shown in Figure 1b, pH levels in both treatment groups rose significantly during the first 6 days of composting, potentially owing to the release of ammonia nitrogen by mineralization [15]. Over time, these pH levels gradually declined as a result of organic acid production and NH_4_^+^ transformation into NH_3_ [23]. When composting was complete, the respective pH values in the samples from groupsCK and T were 8.23 and 8.18, with both meeting the pH requirements for mature compost in the Chinese standard (NY525-2021). In the thermophilic stage, a lower pH was evident in group T as compared to group CK, potentially owing to greater organic and inorganic acid accumulation[25]. These findings indicated that the less basic composting environment in samples from group T was sufficient to inhibit NH_4_^+^ conversion intoNH_3_, thus reducing the loss of nitrogen[26,27].

The germination index (GI) is widely used as a reliable metric to assess the phytotoxicity and maturityof compost[28]. Over the course of composting, the GI in both treatment groups gradually rose with the degradation of volatile fatty acids and other toxic compounds [14], reaching above 80% on day 25 in both treatment groups such that these compost samples were sufficiently mature for use in agricultural settings. When composting was complete, the respective GI values in groups T and CK had risen to 112.4% and 95.7% (Fig. 1c). These data demonstrate that *B.coagulans* X3inoculation can reduce the length of the fermentation cycle, while also improving the maturity of the final compost.

### 3.2 Changes in NH_4_^+^-N,NO_3_^-^-N and TN during composting

No significant changes in concentrations of NH_4_^+^-N were observed during the initial three days of the composting process, whereafter these levels rose, declined, and eventually stabilized(Fig. 2a). As it is influenced by organic nitrogen mineralization and weak nitrification with high temperatures [29], a rapid rise in NH_4_^+^-N concentrations was evident until peaking on day 6 in both treatment groups, with respective maximum concentrations of 621.19, and 571.12 mg/kg in the CK and T treatment groups. These concentrations subsequently fell rapidly from days 6-9, whereafter they declined more slowly. These patterns may be a result of H_3_ volatilization and NH_4_^+^-N transformation into NO_3_^-^-N through the actions of nitrifying microbes[30]. When composting was complete, the levels ofNH_4_^+^-N in both treatment groups were below 400 mg/kg such that all samples met the requirements for maturity[31]. NH_4_^+^-Nlevels in group T were higher than those in group CK, suggesting that *B. coagulans* X3 inoculation resulted in an increase in NH_4_^+^-N content at the maturity stage.

**Fig. 2.**
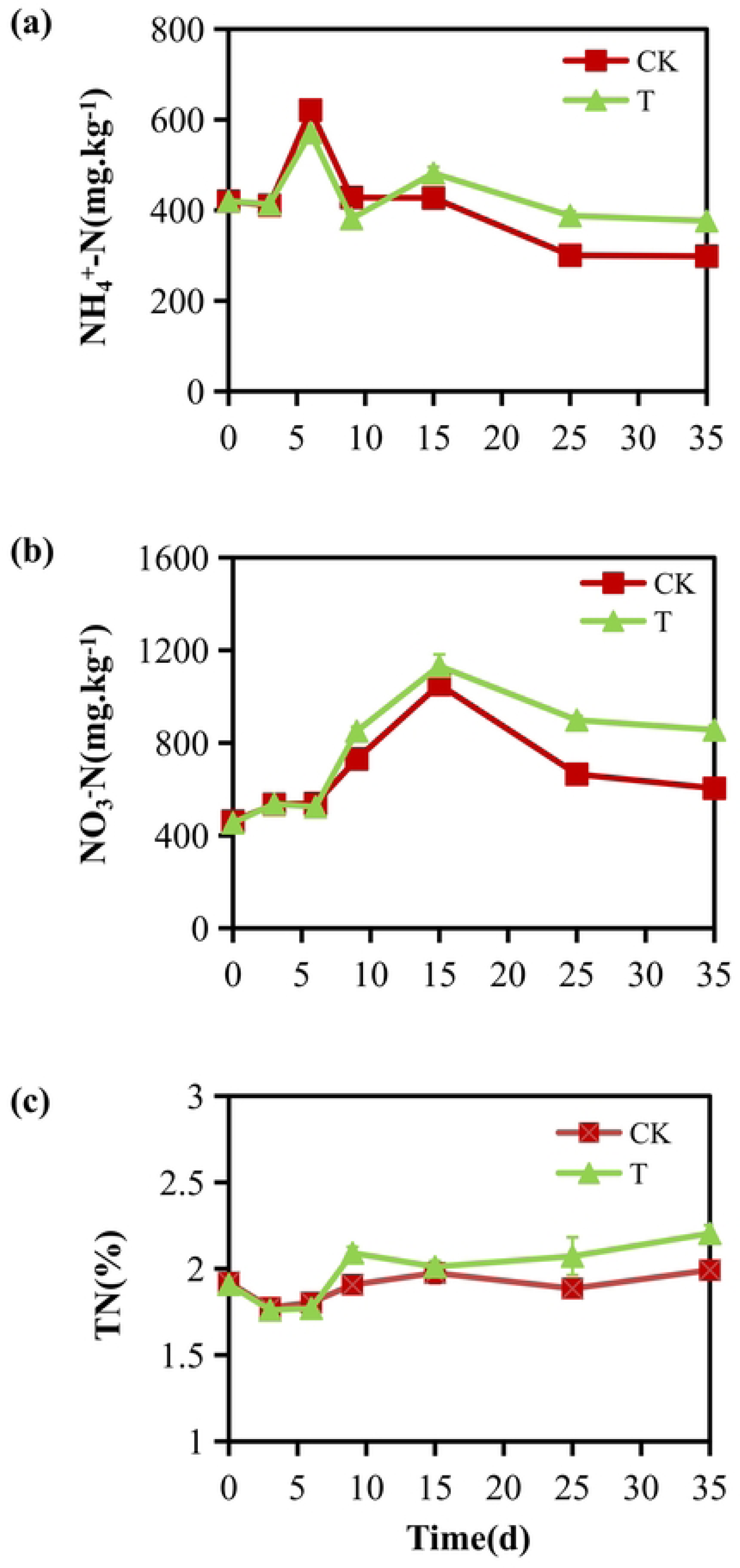
Variations in NH ^+^-N(a), NO ^-^-N(b), and TN(c)levels in the indicated treatment groups over the course of composting(CK, no additive; T, 1% *Bacillus coagulans* X3).

As shown in Figure2b,similar trends inNO_3_^-^-N concentrations were apparent in both groups, initially exhibiting minimal increases during the early stages of the composting process, possibly owing to nitrifyingmicroorganism activity being inhibited by high NH_4_^+^-Nlevels and temperatures [18]. From days 6-15, NO_3_^-^-N levels rose rapidly.Zainudin et al. [32] previously reported that nitrifying activity was enhanced by a composting temperature of approximately 40°C, thereby leading to rapid increases in the concentration of NO_3_^-^-N. Levels in both treatment groups declined until composting was complete, potentially owing to the enhancement of denitrification[33]. When composting was complete, the respective NO_3_^-^-N concentrationsin the CK and T groups were 603.88, and 855.62 mg/kg. This may be attributable to greater active denitrification in group CK, consistent with observed changes in *nirS* gene abundance(Fig. 5).

TN serves as a key index for use when assessing compost quality. TN concentrations in each treatment group trended downward during the first 6 days of composting (Fig. 2c), owing to the high levels of NH_3_emission during the thermophilic stage[29]. After this period, these levels rose gradually owing to the greater loss of carbon relative to nitrogen over the course of composting [33]. A downward trend was observed in both groups as composting temperatures declined, potentially owing to NO, N_2_O, and N_2_ emission via denitrification. When composting was complete, a TN concentration of 2.204% was observed in group T, with this being significantly higher than that in group CK(p<0.05), demonstrating that the addition of *B. coagulans* X3 reduced the loss of nitrogen and increased overall TN content.

### 3.3 Changes in bacterial community diversity

Figure 3a presents the relative phylum level abundance of the bacteria detected in different composting samples. The dominant phyla identified in these analyses were Firmicutes, Proteobacteria, Bacteroidetes, Actinobacteriota, and Chloroflexi, comprising >90% of all sequence reads, in line with findings in organic solid composting systems reported previously[15,22]. Higher levels of relative Firmicutes abundance were evident in the early mesophilic-thermophilic stage as compared to the maturation phase, consistent with the ability of Firmicutes species to generate thermostable endospores capable of surviving under the higher temperatures that arise during composting. Firmicutes are reportedly capable of decomposing cellulose, macromolecular proteins, and other forms of organic matter [34]. On day 6, the relative abundance of Firmicutes in groups T and CK was 50.4% and 28.3%, respectively, with a similar difference also being evidenton day 15(42.2% in T and 19.0% in CK). These data suggest that microbial inoculation stimulated Firmicutes growth in the thermophilic and cooling stages, thereby benefiting the decomposition of organic matter. Proteobacteria, Bacteroidetes,and Actinobacteriota also serve as important mediators of organic matter decomposition and the cycling of carbon, nitrogen, and sulfur [29,35]. Proteobacteria abundance in the CK group rose from 19.5% on day 3 to 31.8% on day 6and 35.7% on day 15, whereas no significant changes in these levels were observed in group T. This suggests that microbial innoculation reduced the abundance of Proteobacteria during the thermophilic-cooling stage.Higher relative levels of Bacteroidetes and Chloroflexi were evident during the maturation stage relative tothe mesophilic-thermophilic stage. In contrast, relative Actinobacteriota abundance was reduced in the cooling-maturation stage relative to the mesophilic-thermophilic stage.

**Fig. 3.**
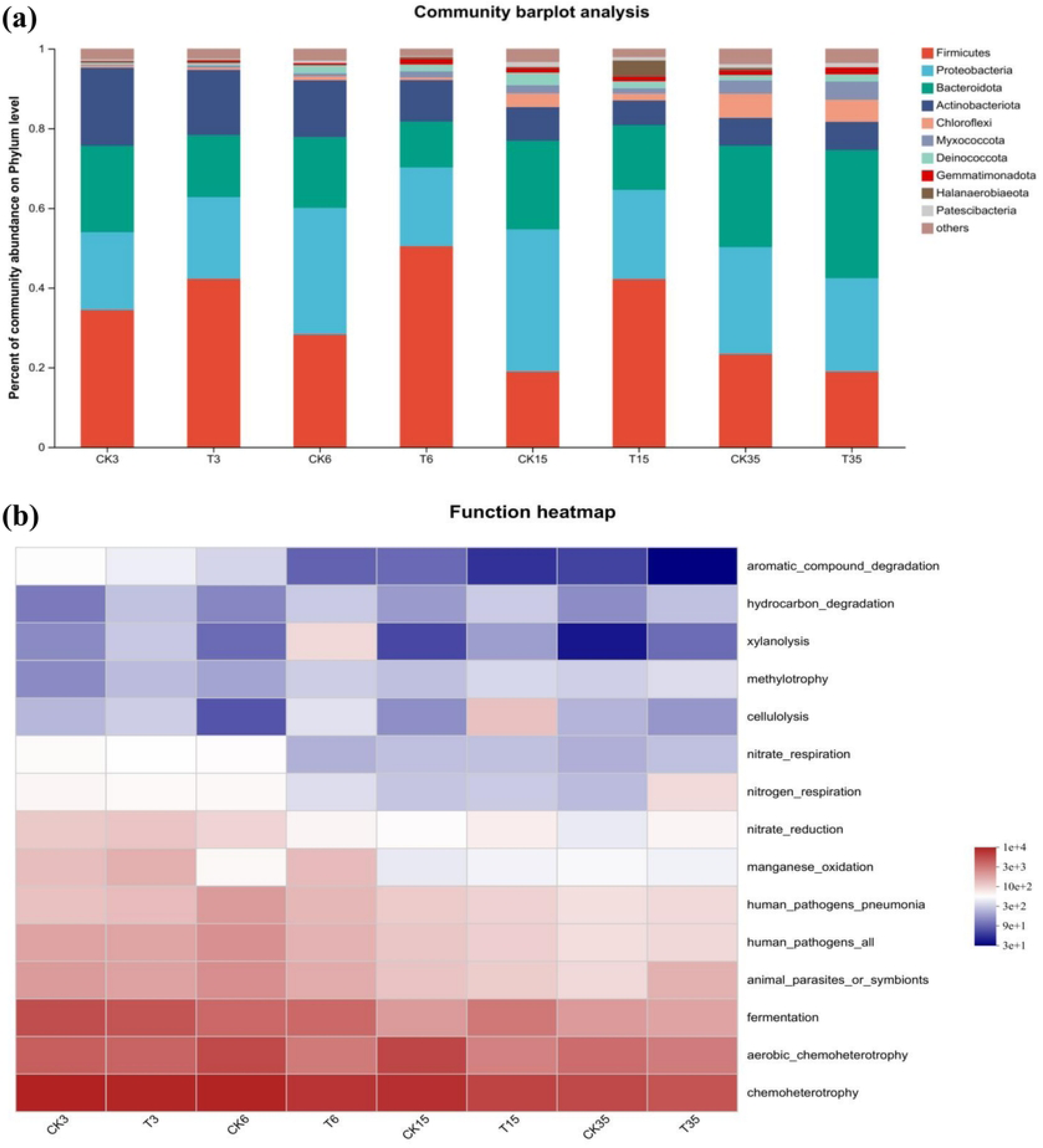
Phylum-level changes in bacterial community composition (a) and predictive analyses of microbial metabolic functionality (b) in different treatment groups over the course of composting(CK, no additive; T, 1% *Bacillus coagulans* X3).

The FAPROTAX database was leveraged to predict microbial functions in the compost samples from both experimental groups, with the top 15 functions being presented in the heatmap in Figure 3b. Relative chemoheterotrophy-related sequence abundance levels were highest, followed by aerobic chemoheterotrophy and fermentation.Relative to the CK group, group T exhibited greater abundance of sequences associated with hydrocarbon degradation, xylanolysis, and cellulolysis during the thermophilic-cooling stage. This difference may be a consequence of greater Firmicutes abundance in the T treatment group. These findings further suggest that *B. coagulans* X3 addition was sufficient to enhance carbohydrate metabolism and to improve the products produced through the composting of cattle manure.

### 3.4 Changes in nitrogen transformation-related functional gene abundance

Changes in *amoA*, *nirS*, *nirK*, and *nosZ* gene abundance have direct or indirect effects on NH_3_, NO, and N_2_O emission, thereby resulting in the loss of nitrogen throughout composting. As demonstrated in Figure 4, the abundance levels of denitrifying genes (*nirS*, *nirK*, and *nosZ*)were higher than those of the nitrifying gene *amoA*, as has been observed in other composting-focused studies[18,22,29]. This suggests higher levels of denitrification activity as compared to nitrification activity in this experimental context.

**Fig. 4.**
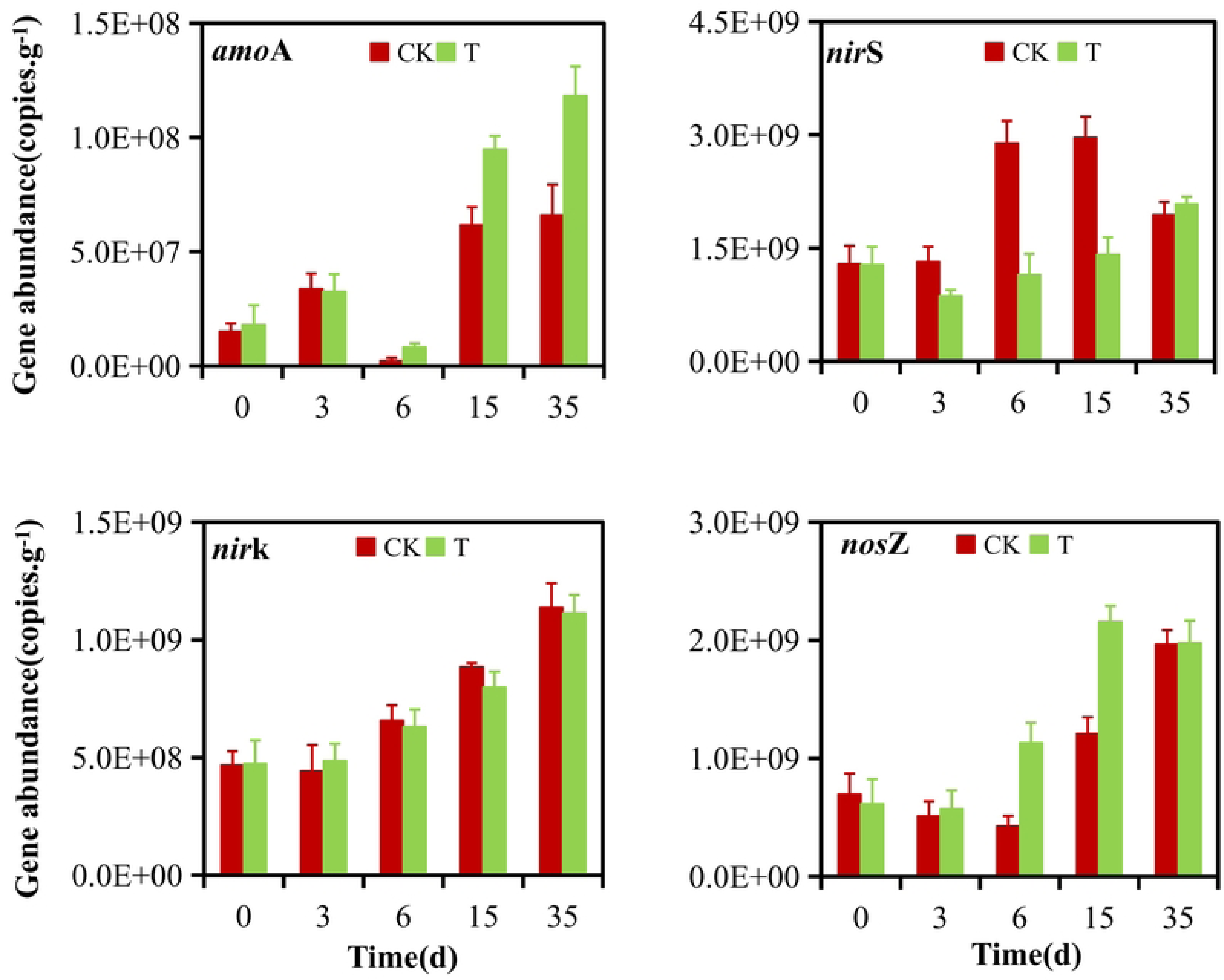
Nitrogen transformation-related functional gene abundance over the course of composting(CK, no additives; T, 1% *Bacillus coagulans* X3).

The enzyme encoded by *amoA*can oxidize ammonia into NH_2_OH, thereby affecting theemission of NH_3_. Both treatment groups exhibited lower *amoA* gene abundance in the early stages of composting relative to the cooling-maturation stage, with the minimum expression being evident on day 6. This may be because higher temperatures inhibit ammonia-oxidizing bacterial growth and activity[36,37]. As temperatures declined,*amoA* copy number rose rapidly during the cooling-maturation stage. Similar findings have also been reported previously[15,38], with *amoA* gene abundance reportedly falling to the lowest levels following a sustained thermophilic period, before ultimately increasing with the entry of the composted materials into the maturation stage. The *amoA* gene abundance levels in group T were significantly higher than those in group CK in the cooling-maturation stage, suggesting that *B. coagulans* X3 inoculation can augment the oxidation of ammonia, thereby decreasing the loss of nitrogen in the form of NH_3_ emissions during the later stages of the composting process.

Denitrifying genes (*nirS*, *nirK*, and *nosZ*) were generally present at higher levels during the cooling-maturation stage as compared to the initial composting stages, consistent with stronger denitrification activity during the later phases of composting. The *nirS* gene abundance levels were higher than those of *nirK* in both groups in this study, in line with previous reports [18,39]. However, Zhang et al.[15]and Guo et al. [40]observed greater *nirK* gene abundance relative to the *nirS* gene when composting agricultural waste. These inconsistent results may be a consequence of substantial differences in composting materials, management practices, and microbial communities. The *nirK* abundance did not differ significantly between groups, whereas *nirS* abundance in group T was lower than that in group CK on days 6 and 15. Given the important role that *nirS* plays in the process of NO_2_ ^-^-N to NO reduction, this suggests that *B. coagulans* X3 has a positive effect on nitrite reduction, thereby reducing the production of NO. Giles et al. [41] found that N_2_O emission levels were primarily reduced through decreases in the production of N_2_O and increases in N_2_O transformation into N_2_. Significantly elevated *nosZ* gene copy numbers were evident in group T relative to group CK on days 6 and 15, further suggesting that adding *B. coagulans* X3 reduces the emission of the greenhouse gas N_2_O.

## 4. Conclusions

In summary, *B.coagulans* X3 inoculation was herein found to be associated with increases in compost temperature and improved compost maturity. Relative Firmicutes abundance rose significantly relative to that observed in the control group during the thermophilic and cooling stages. Bacterial inoculation significantly bolstered *amoA* and *nosZ* gene abundance, whereas *nirS* gene copy number declined in the course of composting, leading to elevated levels of ammonium nitrogen, nitrate nitrogen, and total nitrogen in the final compost product. As such, the addition of *B. coagulans* X3 represents an effective means of minimizing nitrogen loss when composting cattle manure. However, further research will be necessary to clarify the effects of X3 application when composting different types of organic matter.

## Author contribution

All authors contributed to the study conception and design. Material preparation, data collection and analysis were performed by Zhen Wang, Wei Chen, Biao Liu, Minxi Wu,Lijuan Xuand Yongmei Li. The experiments were designed by Hongmei Yin and Zhaohui Guo. The first draft of the manuscript was written by Biao Liu.All authors commented on previous versions of the manuscript.

